# Excitatory and inhibitory synapses form a tight subcellular balance along dendrites that decorrelates over development

**DOI:** 10.1101/2024.04.30.591814

**Authors:** Sally Horton, Vincenzo Mastrolia, Rachel Jackson, Sarah Kemlo, Pedro M. Pereira Machado, Maria Alejandra Carbajal, Robert Hindges, Roland A. Fleck, Paulo Aguiar, Guilherme Neves, Juan Burrone

## Abstract

A balance between excitation and inhibition is crucial for neurotypical brain function. Indeed, disruptions in this relationship are frequently associated with the pathophysiology of neurodevelopmental disorders. Nevertheless, how this balance is established during the dynamic period of neurodevelopment remains unexplored. Using multiple techniques, including *in utero* electroporation, electron microscopy and electrophysiology, we reveal a tight correlation in the distribution of excitatory and inhibitory synapses along dendrites of developing CA1 hippocampal neurons. This balance was present within short dendritic stretches (<20µm), and surprisingly, was most pronounced during early development, sharply declining with maturity. The tight matching between excitation and inhibition was unexpected, as inhibitory synapses lacked an active zone when formed and exhibited compromised evoked release. We propose that inhibitory synapses form as a stabilising scaffold, to counterbalance growing excitation levels. This relationship diminishes over time, suggesting a critical role for a subcellular balance in early neuronal function and circuit formation.

## Introduction

Neurons receive a large number of both excitatory and inhibitory inputs, mostly along their vast somato-dendritic compartment. The balance between the overall levels of excitation and inhibition received by a given neuron in the brain is tightly controlled^1–3^, a feature that is thought to help maintain the stability of neuronal networks. In fact, a dysregulation in the balance between excitation and inhibition has been linked to many neurodevelopmental disorders, including autism, schizophrenia, and epilepsy^4,5^. However, both the emergence of this balance throughout development, as well as the spatial precision with which it is formed, remain less well understood.

Work characterising the distribution of excitatory synapses received by pyramidal neurons in the cortex and hippocampus has uncovered important patterns in the way inputs are arranged along dendrites, with important consequences for dendritic integration^6–10^. In this context, dendrites have been postulated to locally integrate inputs through the clustered distribution of excitatory synapses and the generation of non-linear dendritic events^11–16^. This clustered synaptic arrangement, which is also present during the early periods of synapse formation^17^, endows dendrites with the ability to perform computations in isolation from the rest of the cell^18,19^. The ability of dendrites to integrate excitatory inputs establishing the spatial (and temporal) precision of inhibition along dendrites^20,21^. Do inhibitory inputs match this local excitation at the level of single dendritic branches, perhaps even sub-dendritic domains, or does this balance occur only over the cell as a whole? Recent findings have proposed the existence of a loose subcellular balance in the distribution of synapses along the dendrites of adult cortical neurons^22^, which agree with functional descriptions measured either in hippocampal slices^9^ or in primary neuronal cultures^10^. Furthermore, live imaging of the dynamics of inhibitory and excitatory postsynaptic compartments has shown that remodelling of both synapse types is spatially clustered along dendrites^23, 24^, in agreement with the fact that local potentiation of excitatory inputs also results in a parallel potentiation of inhibitory inputs by dendrite-targeting interneurons^25^, all of which likely contributes to a local balance. However, how this balance emerges during the period of circuit wiring is unknown. Here, we focused on the emergence of synapses throughout development to uncover spatial rules in the arrangement of excitatory and inhibitory synaptic contacts along growing dendrites.

## Results

### Mapping inputs along basal dendrites reveals a local E/I balance

We mapped the distribution of synapses along the basal dendrites of hippocampal CA1 pyramidal neurons by using fibronectin intrabodies to simultaneously label excitatory and inhibitory postsynaptic compartments in the same neuron^26^ (Fig. 1A-D). Specifically, we carried out in utero electroporation of two intrabodies – one raised against the excitatory postsynaptic scaffold protein PSD95 (PSD95-FingR-EGFP) and another to the inhibitory postsynaptic scaffold protein Gephyrin (Geph-FingR-tdT), as well as a cytoplasmic reporter to label dendrite morphology (mRuby2_smFP FLAG)^27^. At P21, following the peak of synapse formation (Fig. 1B-D), we observed puncta for both probes along the entire dendritic tree, that co-localised with different synaptic markers, indicating labelling of bona fide excitatory and inhibitory synapses (Fig. S1). Importantly, these probes do not interfere with synapse structure or function^26^ and their labelling intensity reflects synapse size (Fig. S1 A-D). As expected, the density of Geph-FingR-tdT puncta along a dendrite was quite low (0.12 synapses.μm-1, SEM= 0.01, n=21 dendrites from 7 cells) and occurred mainly on the dendritic shaft (∼95% on shaft), whereas PSD95-FingR-EGFP puncta were much more abundant (0.8 synapses.μm-1, SEM=0.07, n=21 dendrites from 7 cells) and mostly on dendritic spines (∼80%). Since the levels of excitation and inhibition are thought to be broadly balanced in adult neurons at both a cell-wide and subcellular level^1,22,28–30^, we sought to explore this relationship in more detail.

**Figure 1.**
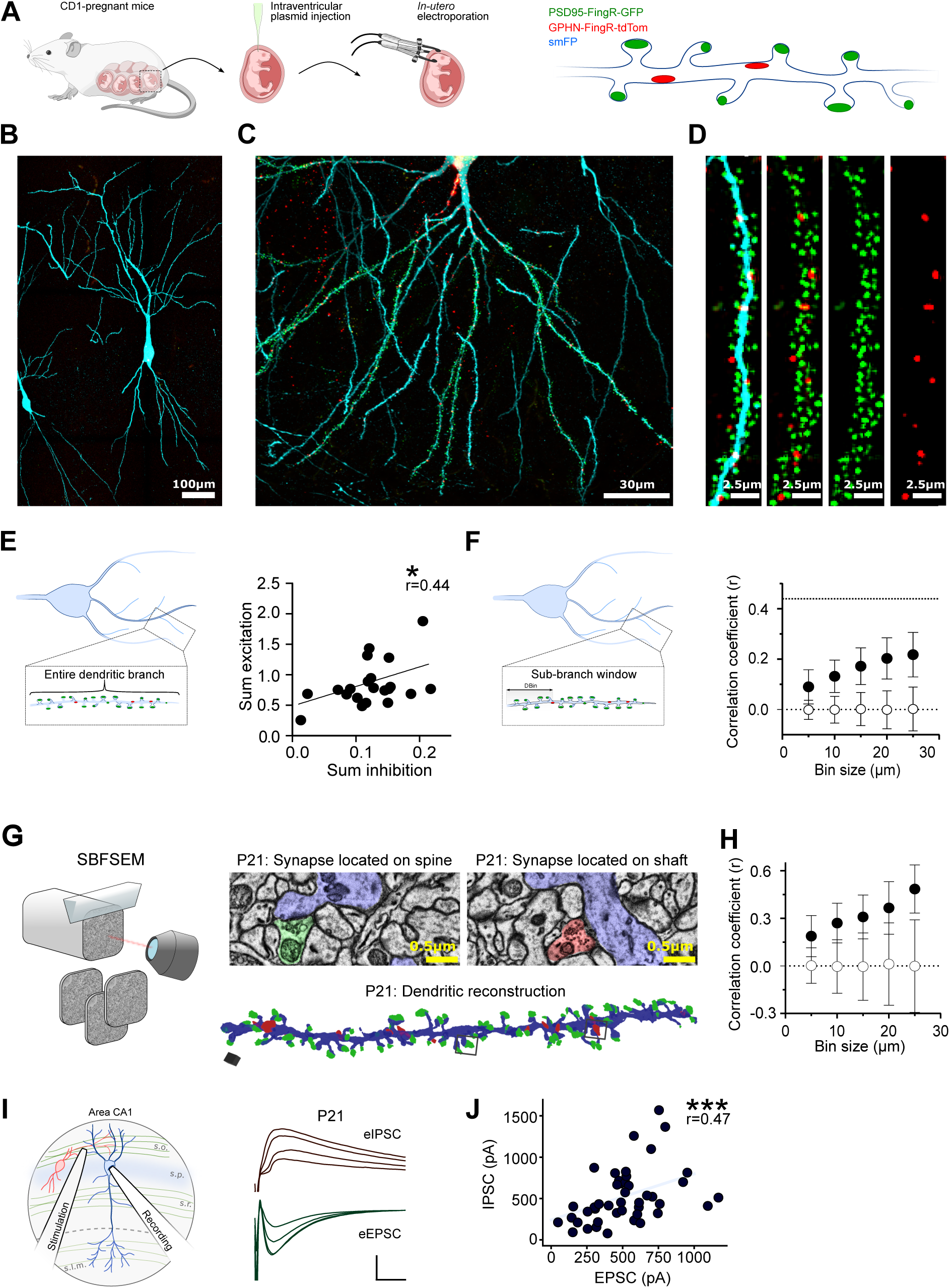
Mapping excitatory and inhibitory synapses along the basal dendrites of CA1 pyramidal neurons at P21, using FingRs, SBFSEM and electrophysiology. (A) Schematic showing delivery of PSD95-FingR-GFP, Gephyrin-FingR-tdTom and smFP into pyramidal neurons by *in utero* electroporation to simultaneously label excitatory (green) and inhibitory synapses (red), as well as the entire neuron (blue). (B-D) Example images of pyramidal neurons in CA1 of a P21 hippocampus at different levels of spatial resolution, from an entire cell (B) to basal dendrites (C) and a single dendrite (D) where synapses are clearly visible. (E) The balance between inhibition and excitation was assessed along the lengths of entire basal dendritic branches. The graph shows the correlation between the sum of excitation as a function of the sum of inhibition at P21. A significant correlation was observed (n=21 dendrites, r=0.4390, P=0.046). (F) The balance between excitation and inhibition was assessed within sub-branch windows (DBin) of different sizes by random sampling different dendritic stretches across the dendrite. Analysis was repeated 10000 times and mean±SD of the correlation coefficients are shown (closed circles). The same analysis was repeated with modelled dendrites where synapses were randomly placed (open symbols). The correlation coefficient increases as a function of Bin Size. Lines are drawn both at 0 and at the correlation coefficient observed when entire branches were analysed (E). (G) Serial block-face scanning electron microscopy (SBFSEM) was carried out in the *stratum oriens* of the CA1 hippocampus from P21 brains. Example single plane images (top, centre) of excitatory (green) and inhibitory (red) synapses formed onto a dendrite (blue). Left image shows an excitatory synapse onto a spine. Right image shows an inhibitory shaft synapse. Below is a 3D reconstruction of a dendrite (blue) showing excitatory (green) and inhibitory (red) synapses. (H) Correlation coefficients increase with bin size in the experimental (closed circles) but not the simulated branches (open circles). (I) Schematic showing the electrophysiology recording and stimulation setup (left) and example traces (right) for evoked excitatory (green) and inhibitory (red) postsynaptic currents measured in the same cell at P21, in response to stimulation with 4 different (increasing) current intensities. (J) A graph showing the correlation between the amplitude of excitatory (EPSC) and inhibitory (IPSC) postsynaptic currents at P21. A significant positive correlation was observed between excitatory and inhibitory currents at P21 (n=11, r=0.4744, P<0.001).

At this peri-adolescent stage, we found that the cumulative fluorescence intensity of excitatory and inhibitory synapses (‘Sum’, Fig. 1E), but not the density (Fig. S2), showed a weak positive correlation. The sum incorporates an indirect assessment of synapse strength and is therefore likely to represent more accurately the functional correlation between excitation and inhibition (see Methods). Bootstrapping analysis confirms this correlation was not present when the same analysis was performed in simulated branches with randomly placed synapses (Fig. S2D). We were also keen to investigate if the levels of excitatory synaptic input predicted the inhibition received along a range of spatial window sizes (bins) within a branch (Fig. 1F). This is because dendrites can integrate local excitatory inputs and initiate non-linear depolarizations, so mapping the fine scale spatial distribution of excitatory and inhibitory synapses is fundamental to understanding dendritic computation^31,32^. Furthermore, excitatory inputs encoding similar features in the environment form clusters along short stretches of dendrites (<20µm), both in young adults and developing neurons^11–13,17,19^. We found that the correlation coefficient between excitation and inhibition increased with bin size, becoming progressively distinct from randomly placed synapses and suggesting correlations were present at a sub-dendritic level that match the spatial domains of functional clusters (Fig. 1F). To confirm the identified subcellular correlation, we performed serial block-face scanning electron microscopy (SBFSEM) to map excitatory and inhibitory synapses along the basal dendrites of P21 CA1 hippocampal neurons (Fig. 1G, 392 synapses (338 on spines and 54 on shafts) over 5 dendrites covering a length of 262µm). Again, correlations between the summed excitation and inhibition were found along short stretches of dendrite (Fig. 1H). To provide a functional readout of this balance, we conducted whole-cell recordings of pyramidal neurons in P21 acute hippocampal slices and measured evoked currents for both excitatory and inhibitory inputs onto basal dendrites by placing a stimulating electrode in the stratum oriens (Fig. 1I). By varying the strength of stimulation to activate a range of inputs locally, we found that EPSC and IPSC amplitudes were correlated, in agreement with previous findings^28^ (Fig. 1J). Altogether, our data shows that at P21 there is a loose subcellular balance between the level of excitation and inhibition along dendrites, which extends to short dendritic domains containing only a small cluster of synapses.

### A subcellular E/I balance is highest during early development

To understand whether the local balance between excitatory and inhibitory synaptic inputs is a feature that emerges during development or is present from the moment of synapse formation, we characterised the distribution of synapses along dendrites during the peak of synapse formation, from P7 to P14 (Fig. 2A-C). We exploited the ability of fibronectin intrabodies to label postsynaptic compartments that lack any clear morphological characteristics, which is particularly important for synapses that form directly onto the dendritic shaft, such as inhibitory synapses, and excitatory synapses during early in development. We found that whereas the density of GABAergic synapses was similar throughout development, excitatory synapse density increased dramatically, particularly from P14 to P21 (Fig. 2D-E). In parallel, we saw a gradual transition in the location of excitatory synapses from the dendritic shaft at P7, to dendritic spines by P21 (Fig. 2G). Inhibitory synapses remained primarily on the shaft throughout development (Fig. 2F). Importantly, at P7, the PSD95-FingR-EGFP puncta, including those on the dendritic shaft, co-localised with the presynaptic marker Bassoon and the Geph-FingR-tdT puncta co-localised with presynaptic vGat, indicating these were bona fide synapses (Fig. S1G-H). Surprisingly, at P7, before the bulk of excitatory synapses form, the levels of excitation and inhibition were tightly balanced across individual dendrites, both at the level of synapse density (Fig. S2A) as well as the sum of synaptic inputs (Fig. 2H). In fact, the level of correlation remained high until P14 but was substantially reduced by P21 (Fig. 2I). We further explored the spatial extent of this correlation using the spatial binning analysis (Fig. 2J-K, Fig. S2B). Once more, sub-dendritic correlations emerge within short spatial bins. The reduction in correlation between excitation and inhibition across development prompted us to look at the distribution of each synapse type independently. The distance between neighbouring inhibitory synapses deviated from a random distribution across all ages, whereas excitatory synapses gradually became more randomly distributed, so that their distribution at P21 could be well described by an exponential function (Fig. S2C). This suggests synapse formation starts as a coordinated process, while at more mature stages excitatory synapse development becomes increasingly decorrelated. Interestingly, a tight correlation between excitation and inhibition was also observed in dissociated hippocampal neurons (Fig. S3)^33^, suggesting that this synaptic distribution does not require a precise circuit architecture but, instead, likely reflects local dendritic rules.

**Figure 2:**
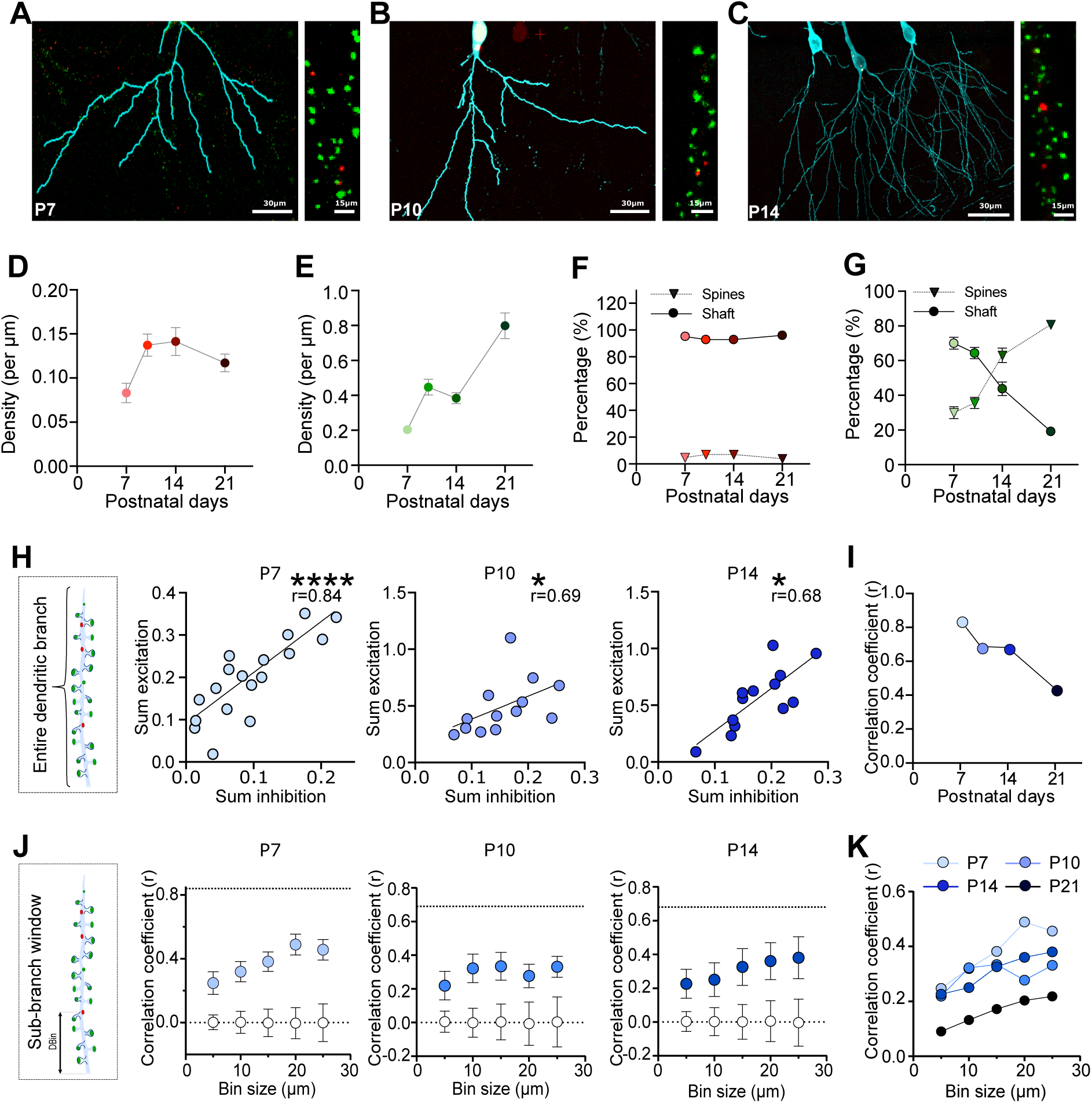
Excitation and inhibition are balanced at the subcellular level in early development. (A-C) Example images of basal dendrites of pyramidal neurons imaged in fixed brains expressing PSD95-FingR-GFP (green), Gephyrin-FingR-tdTom (red) and smFP (blue) at P7 (A), P10 (B) and P14 (C). (D-E) A plot of the density of inhibitory (D) and excitatory (E) synapses across postnatal days (darker colours correlate with age). (F-G) A plot of the location (either on the dendritic shaft or a spine) of inhibitory (F) and excitatory (G) synapses as a function of postnatal age. (H) Schematic showing the spatial domain (entire dendrite) of the correlation analysis carried out and graphs showing the correlation between the sum of excitation as a function of the sum of inhibition from P7 (far left graph) to P14 (far right graph). A tight and significant positive correlation was observed at P7 (n=18, r=0.8431, P<0.0001), P10 (n=13, r=0.6868, P=0.0118), and P14 (n=13, r=0.6813, P=0.01). (I) A summary plot of the correlation coefficients (r) measured in (H) as a function of postnatal days. The correlation coefficient is highest a P7 and gradually decreases throughout development. (J) Schematic showing how the balance between excitation and inhibition was assessed within sub-branch windows (DBin) of different lengths. Graphs showing the correlation coefficient as a function of Bin Size. Correlations increased with bin size in experimental (closed circles) but not simulated (open) branches at all analysed ages. (K) A summary plot showing an overlay of the data shown in Fig 2J and Fig 1F showing lower levels of correlation at P21.

### The emergence of a functional E/I balance during development

Next, we measured evoked currents for both excitatory and inhibitory inputs through development (Fig. 3A-B). We observed a correlation between EPSC and IPSC amplitude across all ages that peaked at P14 (Fig. 3A-B). Although we observed a decline in both structural and functional correlations between P14 and P21, at earlier stages the weaker functional correlation was at odds with that observed with intrabodies. A noticeable feature of IPSCs at P7 was their small amplitude (Fig. 3C), suggesting immature levels of evoked inhibition may be the reason for this structure-function misalignment. In contrast, we found no changes in the amplitude or frequency of spontaneous IPSCs (sIPSCs) throughout development (from P7 to P21) (Fig. 3D-E).

**Figure 3.**
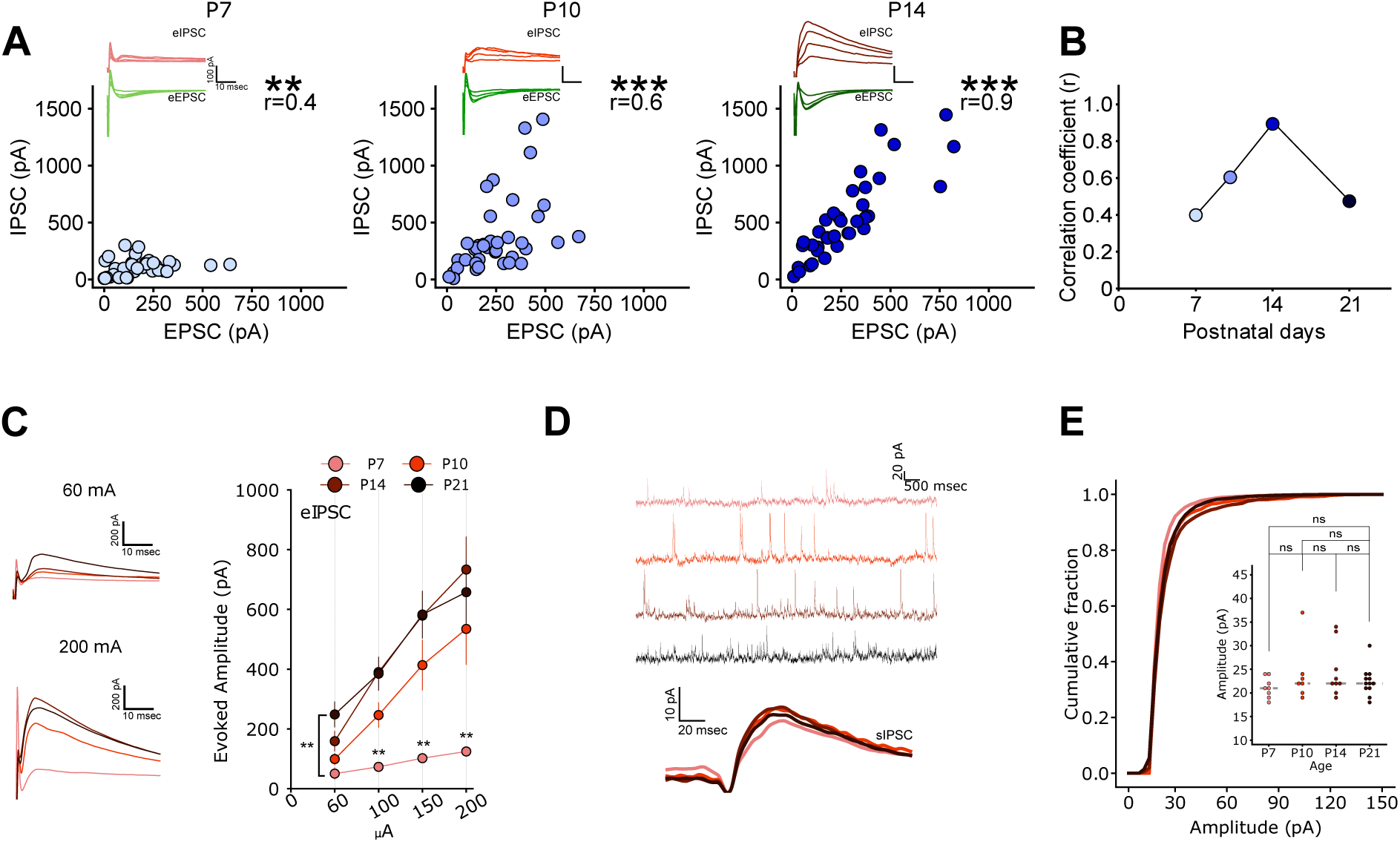
There is a functional balance between excitation and inhibition at the subcellular level in early development. (A) Graphs showing the correlation between the amplitude of excitatory (EPSC) and inhibitory (IPSC) postsynaptic currents, from P7 (far left graph) to P14 (far right graph). A significant positive correlation was observed between excitatory and inhibitory currents at P7 (n=10, r=0.3990, P<0.001), P10 (n=10, r=0.6038, P<0.001) and P14 (n=11, r=0.8803, P<0.001). Example EPSC (green) and IPSC current traces are shown above each graph. (B) A summary plot of the correlation coefficients (r) measured in (A) as a function of postnatal days. Note the increase in correlation from P7 to P14 and the drastic decrease at P21. (C) Evoked inhibitory postsynaptic currents (IPSCs) recorded in CA1 pyramidal neurons from P7 to P21. Notice the weak IPSCs measured at P7. (D) Example traces of spontaneous IPSCs (sIPSCs) recorded in CA1 pyramidal neurons from P7 to P21. (E) A graph showing no change in the amplitude of sIPSCs recorded throughout development (from P7 to P21). Inset shows mean sIPSC amplitudes across ages.

### A switch in inhibitory synapse composition between P7 and P10

To understand the inability of immature inhibitory synapses to undergo evoked release reliably, we characterised their structural development in more detail. Super-resolution imaging of vGat-labelled presynaptic boutons using dSTORM (Fig. 4A) revealed no significant differences in the volume of vGat puncta across all ages (Fig. 4B), confirming that inhibitory synapses do not dramatically change during development. To further characterize inhibitory synapses, we stained for the ubiquitous presynaptic active zone (AZ) marker Bassoon (Fig. 4C), which is usually present in all boutons, regardless of identity. We found that whilst the majority (∼97%) of PSD95-FingR-EGFP puncta were associated with Bassoon (Fig. S1G), only a minority (37%) of all Geph-FingR-tdT puncta did so at P7 (Fig. 4C, top row). This contrasts with the ∼97% co-localisation of Geph-FingR-tdT and vGat (Fig. S1H), suggesting that a presynaptic compartment was present, but lacked Bassoon. Indeed, only ∼40% of endogenous vGat puncta contain either Bassoon (Fig. 4C middle row), or RIM1/2 another essential AZ protein (Fig. 4C bottom row). By P10, the vast majority (∼90%) of vGat positive synapses are clearly labelled for both AZ markers and remained highly co-localised at P21 (Fig. 4C). These results suggest that at P7, inhibitory synapses are immature and do not yet have a fully formed AZ, a feature that could explain the small evoked IPSCs (Fig. 3C).

**Figure 4.**
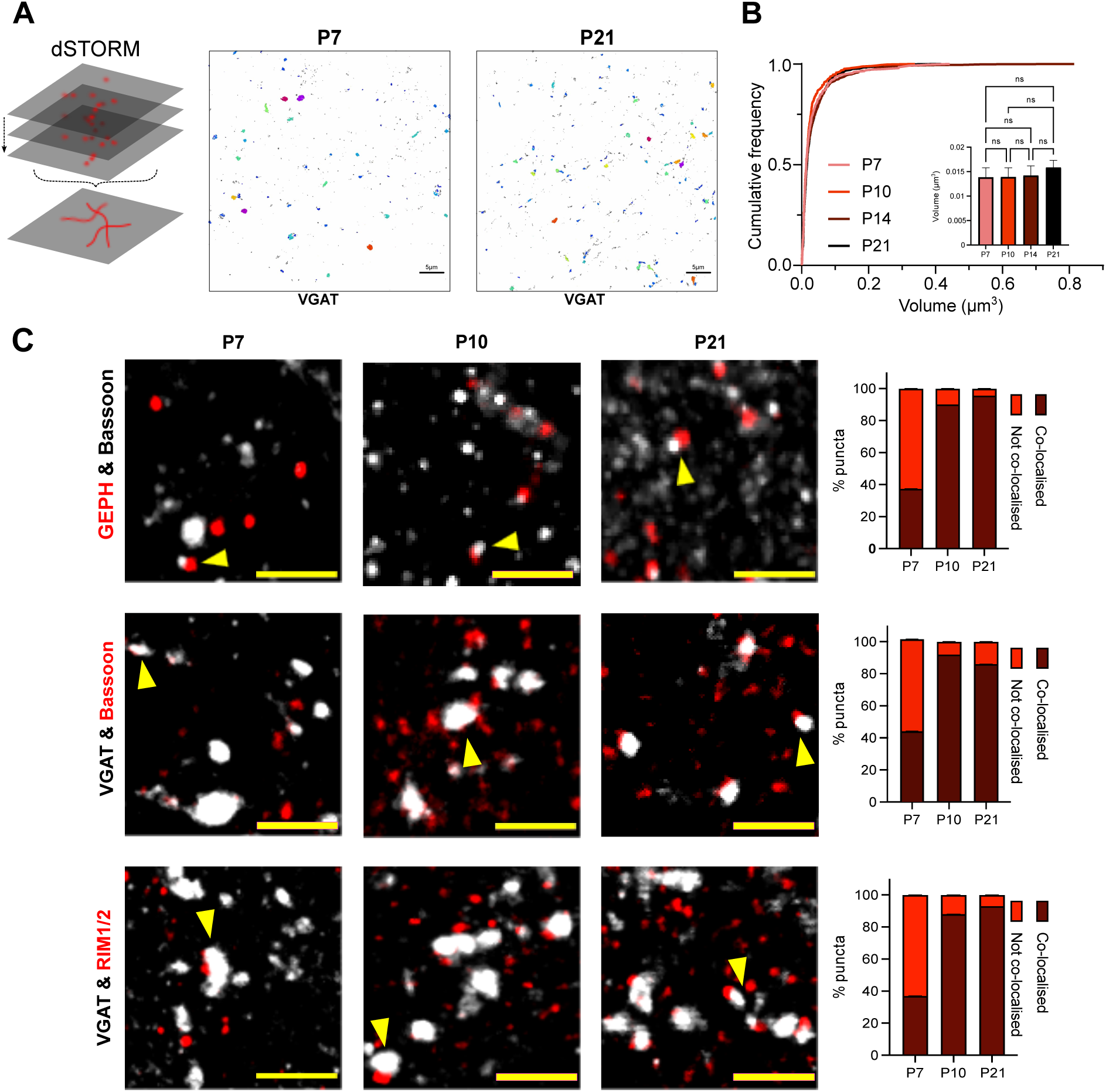
Inhibitory synapses provide a structural scaffold early in development. (A) Schematic showing the principles of direct Stochastic Optical Reconstruction Microscopy (dSTORM) used to image VGAT in fixed brain slices. Clusters of VGAT localizations were identified and volumes measured using Voronoi diagrams in Point Clouds Analyst (PoCA) shown in the example images (P7 left and P21 right). (B) Plot of the cumulative distribution of VGAT cluster volumes showing no difference across age, from P7 to P21, darker red lines represent older synapses, (P7 n=394 clusters, P10 n=417 clusters, P14 n=797 clusters, P21 n=690 clusters, images taken from at least 3 slices from at least 3 separate brains). Inset shows median VGAT volume +/-95% CI, Kruskal-Wallis test with Dunn’s multiple comparisons (ns = p>0.05). (C) Confocal imaging of inhibitory synapses across development (P7, P10 and P21). Top row: example images of Geph-FingR-tdT puncta (red) co-stained with antibody against Bassoon (white). Example images of synapses co-stained for vGAT (white) and Bassoon (red). Bottom row: example images of synapse stained for vGAT (white) and Rim 1/2 (red). Bar graphs (far right) indicate proportions of synapses that showed clear co-localisation between the two markers for each row, across different postnatal days. Only a minority of GEPH-FingR-tdTom puncta were colocalised with bassoon at P7 (37%, n=99, SEM=0.05%), but this colocalization was 90% at P10 (n=132, SEM=0.03%), and 95% at P21 (n=160, SEM=0.02%). Similarly, at P7, the majority of VGAT positive synapses did not colocalized with the active zone markers bassoon (43%, n=99, SEM=0.05%) or RIM1/2 (37%, n=100, SEM=0.05%). At P10, the majority of VGAT puncta colocalised with bassoon (92%, n=100, SEM=0.03%) and RIM1/2 (88%, n=100, SEM=0.03%), which was also the case at P21 (bassoon 86%, n=100, SEM=0.03; RIM1/2 93%, n=100, SEM=0.03%).

Together, our data shows that whereas excitatory synapses on basal dendrites are fully functional by P7, inhibitory synapses initially form with an immature AZ and are not fully competent for evoked release. These synapses can, however, release GABA spontaneously (Fig.3D-E), in agreement with previous descriptions of synapses lacking the active zone protein RIM1a^34,35^. Mechanistically, our findings show that despite the lack of evoked release, inhibitory synapses form structural contacts with postsynaptic strengths that distribute along dendrites in a way that matches excitation. More interestingly, dendritic inhibition provides only a weak functional balance to excitation during the early period of brain development, with synapses remaining latent until they are plugged in to the network by the acquisition of an active zone and the ability to carry out evoked neurotransmitter release. Our data suggests synapses form through spatially patterned inputs that set up an exquisite level of crosstalk between excitation and inhibition, which is likely to play a role in allowing circuits to develop in a stable manner.

## Discussion

We explored the spatial rules that underlie the distribution of synapses along dendrites during circuit wiring. Using both genetically encoded fluorescent reporters to label synapses, as well as 3D EM reconstructions, we mapped the location and properties of synapses along the basal dendrites of CA1 hippocampal neurons. We found that excitatory and inhibitory synapses are correlated along dendrites across all developmental time-points (P7 to P21), although the correlation was highest during the earlier periods of synapse formation, particularly at P7, and sharply declined during peri-adolescence. By exploring the arrangement of synapses along dendrites in more detail, we uncovered a non-random distribution of inhibitory synapses that explains the tight correlations observed here. Functional measures of excitation and inhibition also showed a correlation, although it was generally weaker at earlier time-points, such as at P7. This structure-function discrepancy is explained by the fact that inhibitory synapses initially form onto basal dendrites as immature compartments that lack an active zone and the ability to reliably carry out evoked release of GABA. We propose that inhibitory synapses provide a structural dendritic scaffold that allows the formation of excitatory synapses to take place in a balanced manner.

The notion of a local dendritic balance between excitation and inhibition was originally described in elegant experiments carried out in primary neuronal cultures^33^, which we confirmed here with our synaptic probes. More recently, a broad and detailed map of excitatory and inhibitory synapses uncovered a loose subcellular correlation across the dendrites of adult cortical layer 2/3 pyramidal neurons^22^. At the whole dendrite level, the magnitude of the correlations were very similar to those reported here for P21 CA1 pyramidal neurons (Fig. 1) and suggests common mechanisms may be at play. Together with the tight correlation observed in dissociated hippocampal neurons (Fig. S3), these findings suggest that the rules for the generation of balanced inputs may be independent of brain region (cortical versus hippocampal) or circuit structure (intact tissue versus dissociated cultures). In other words, they may represent a universal property in the way synapses are formed onto the dendrites of pyramidal neurons. These results also tally well with recent experiments showing that sparse stimulation of presynaptic CA3 neurons resulted in balanced postsynaptic excitatory and inhibitory currents measured in adult CA1 neurons^28^. How this feedforward inhibition is wired up to achieve such a balance remains unknown but may arise, at least in part, from the patterned arrangement of synaptic inputs described here.

Dendrites have a unique ability to integrate excitatory inputs locally, resulting in dendritic non-linearities that enhance the information capacity of neurons^19,36,37^. The basal dendrites of pyramidal neurons in both the cortex and hippocampus, for example, show clear supra-linear events when sufficient synapses are activated^38–41^. In vivo imaging of synaptic events has shown that inputs that code for similar features in the environment form clusters along 10-20µm stretches of dendrite, providing the synaptic basis for the generation of non-linear dendritic events^11,12,14^ and similar clusters were observed when mapping the connectivity of CA3 axons onto CA1 dendrites^15^. Although we did not map the functional clustering of synaptic inputs onto CA1 neurons, our results suggest that inhibition is well placed to provide a local balance to ongoing excitation with a precision that matches the extent of clustered inputs (Fig. 1-2). A proper functional characterisation of excitatory and inhibitory events along dendritic segments remains an important experiment for the future and will likely require sensitive probes with improved spatial and temporal resolution^42^.

Our main finding is the description of a strong correlation between excitation and inhibition during the early period of development, which suggests that the process of synapse formation takes place in a non-random manner. Interestingly, clustered excitatory inputs have been shown to emerge very early on (from P7 onwards)^31,43^, which matches the developmental period over which inputs are highly correlated (Fig. 2). Although the mechanisms for forming correlated inputs remain a mystery, recent findings may provide clues. For example, uncaging GABA onto young, P7, dendrites resulted in the formation of both a gephyrin cluster and the emergence of a new dendritic spine close by^44^. Conversely, repeated stimulation of a dendritic spine in the vicinity of an inhibitory axon, elicited the release of endocannabinoids from the postsynaptic dendrite, the growth of a presynaptic bouton in the neighbouring axon and the formation of a GABAergic synapse^45^. Together, these coordinated synaptogenic events that rely on the local cross-talk between excitatory and inhibitory synapses could explain the high levels of correlation observed here. In addition, local plasticity rules may also foster a correlation. Induction of NMDAR-dependent LTP of excitatory synapses on the distal dendrites of cortical pyramidal neurons also resulted in a parallel potentiation of dendritic-targeting inhibitory inputs^25^. In line with this, ongoing turnover of synaptic inputs appear to take place in clusters of both excitatory and inhibitory synapses, suggesting that plasticity is spatially coordinated at the dendritic level^23, 24^. Overall, it appears that excitation and inhibition are not only formed in a coordinated manner, but also follow plasticity rules that tend to keep this local balance intact.

Finally, we find that inhibitory synapses are initially formed in an immature state, lacking an AZ and evoked release (Fig. 3-4). Previous work has shown the time-course of synapse formation-the coming together of a presynaptic bouton and a postsynaptic compartment-takes place over a few hours for both excitatory^46–48^ and inhibitory synapses^49,50^. However, the maturation of these newly formed synapses may take longer, perhaps even days^51,52^. To date, little is known about the assembly of the presynaptic AZ in GABAergic boutons and its role in controlling the functional maturation of inhibition during development. A recent study measured the increase in functional inhibition in the hippocampus during the first two postnatal weeks, that peaks at P10, and promotes the appearance of hippocampal ripples^53^. In line with this study, we provide evidence in intact tissue that inhibitory synapses are able to form pre-and post-synaptic contacts that contain vGAT and gephyrin but lack key AZ proteins, such as RIM1/2 and Bassoon, at P7 but are present by P10 (Fig. 4) when evoked release was observed (Fig. 3). This endows inhibitory synapses the ability to establish a foothold on dendrites, without having a functional impact on dendritic integration, at least until the AZ is formed. Although it remains unclear why this latency period exists, it is tempting to speculate that it may provide a temporal buffer to allow inhibitory synapses to transition from their early excitatory role to one of inhibition later on, an event that is thought to take place within the first postnatal week^44,54^. Regardless of the reason for this developmental trajectory, we provide evidence that inhibitory synapses follow a different mechanism of synapse assembly to that observed for excitatory synapses. More importantly, we propose that inhibitory synapses are built following a non-random dendritic layout that dictates the balance between excitation and inhibition along developing dendrites.

## Acknowledgments

We would like to thank Laura Andreae and Matthew Grubb for useful feedback and comments on the manuscript. We would also like to thank Nicolas Bourg and Abbelight for the SAFeRedSTORM module and software, as well as Olympus for the IX3 microscope used for dSTORM and Benjamin Compans for help with both using the platform as well as with dSTORM techniques in general. This research was funded in whole, or in part, by the Wellcome Trust (215508/Z/19/Z to JB and an equipment grant 108461/Z/15/Z to RF and JB). For the purpose of open access, the author has applied a CC BY public copyright licence to any Author Accepted Manuscript version arising from this submission. This work was also supported by a BBSRC project grant (BB/S000526/1) to JB. SH was funded by an MRC-ITND PhD studentship.

## Author contributions

G.N. and J.B. conceived the project. S.H., V.M., R.J., G.N., and J.B. developed and designed the experiments. S.H. undertook the in utero electroporation experiments, immunolabelling, confocal imaging and analysed the confocal data. G.N., and S.H segmented and analysed the SBFSEM datasets. P.M. P.M., M.A.C., and R.A.F. provided technical expertise and guidance on EM techniques. P.M.P.M., and M.A.C. processed the samples for SBFSEM and acquired the images. V.M. undertook the electrophysiology experiments and analysis. R.J. carried out the STORM imaging and the data analysis. G.N. wrote the IgorPro analysis scripts. S.H., V.M., R.J., G.N., and J.B. wrote the paper with feedback from authors. PA performed data modelling. SK characterised the tools used (FingRs).

## Materials and Methods

### Acute Slice Preparation

CD1 mice (age P7, P10, P14 and P21) were anaesthetised with a cocktail of Ketamine (200mg/kg) and Xylazine (40mg/kg) and decapitated. Brains were quickly harvested and kept in cold slicing solution (93 mM NMDG, 20 mM Glucose, 20 mM Hepes, 2.5 mM KCl, 30 mM NaHCO2, 1.2 mM NaH2PO4, 7 mM Na-Ascorbate, 2 mM Thiourea, 4 mM Na-Pyruvate, 0.5 mM CaCl2, 10 mM MgCl2, pH 7.35 carbonated with 95% O2/5% CO2) for 1-2 minutes. Brains were transferred on a vibratome (7000smz-2, Campden Instruments, UK) and sagittal dorsal hippocampal slices (300 mm thickness, speed 0.05 mm/s, 70 Hz/1 mm amplitudes) were collected and incubated for 20 minutes in carbonated slicing solution at 32° C. Slices were then transferred in holding solution (95 mM NaCl, 12 mM Glucose, 20 mM Hepes, 2.5 mM KCl, 30 mM NaHCO2, 1.2 mM NaH2PO4, 7 mM Na-Ascorbate, 2 mM Thiourea, 4 mM Na-Pyruvate, 2 mM CaCl2, 2 mM MgCl2, pH 7.35 carbonated with 95% O2/5% CO2) at room temperature and let recover for at least 2 hours.

### Electrophysiology

After recovery, dorsal hippocampal acute slices were transferred to a recording chamber equipped with a custom-built microscope (Cosys Ltd) supplied with Dodt gradient contrast and infrared light and superfused with recording aCSF solution (124 mM NaCl, 5 mM KCl, 1.25 NaH2PO4, 6 mM Glucose, 26 mM NaHCO3, 5 mM Hepes, 2 mM CaCl2, 1 mM MgCl2, pH 7.35 carbonated with 95% O2/5% CO2) at 32° C. Spontaneous and evoked currents were recorded with a Multiclamp 700B amplifier (Axon, Molecular Devices) and digitised with the Digidata 1550B digitizer (Axon, Molecular Devices) with 10 kHz sampling rate and 5 kHz low-pass filter. Data was acquired with the software Clampex 10.7 (Axon, Molecular Devices). For spontaneous postsynaptic currents recordings, a CA1 pyramidal neuron was selected and whole-cell configuration was achieved using a borosilicate pipette (World Precision Instruments) with a resistance of 4-5 MW and containing internal solution (135 mM KMeSO4, 5 mM KCl, 10 mM Hepes, 0.2 mM EGTA, 5 mM MgATP, 0.3 mM NaGTP, 10 mM Na2-Phosphocreatinine, 1 mM MgCl2, 5 mM QX-314, pH 7.3). The patched neuron was either held at-70 mV or at 0 mV, to record spontaneous excitatory postsynaptic currents (sEPSCs) or spontaneous inhibitory postsynaptic currents (sIPSCs), respectively. For evoked postsynaptic currents recordings, a stimulating electrode connected to an external stimulator (A.M.P.I., Israel) and filled with recording aCSF was positioned in the centre of the stratum oriens in the CA1 area of the hippocampus. Approximately 50 mm more distally, a CA1 pyramidal neuron was selected and whole-cell configuration was achieved using a borosilicate pipette with a resistance of 4-5 MW and containing internal solution. 2 pulses of current (0.2 msec pulse duration, 20 Hz) were delivered by the stimulating electrode at 4 current intensities (60, 100, 150, 200mA) to trigger the presynaptic terminals targeting the patched neuron. During the stimulation, the patched neuron was either held at-70 mV or at 0 mV, to record evoked excitatory postsynaptic currents (eEPSCs) or evoked inhibitory postsynaptic currents (eIPSCs), respectively. All data was analysed and plotted using a custom-made Python script.

### In utero electroporation

Prior to IUE, DNA was prepared to a concentration of 1-2µg/µl, and fast green dye was added at a concentration of 0.3mg/ml. In utero electroporation (IUE) was undertaken on CD1 mice at E14.5. Dams were subcutaneously injected with analgesic twenty minutes prior to the IUE surgery (0.1mg/Kg Vetagesic). Anaesthesia was induced by 5% isoflurane (1.2ml/min oxygen) and maintained using 2% isoflurane. IUE was performed as described in published protocols (Saito and Nakatsuji, 2001 and Szczurkowska et al. 2016). DNA was injected into the lateral ventricles via surgical needles pulled from glass thin-walled capillaries (World Precision Instruments, TW15-OF4). A triple-electrode configuration was used to apply 5 electrical pulses: 50ms duration, 30V, 950ms interval (Nepagene, CUY21-EDIT). Embryos were returned to the abdominal cavity, and the dam’s muscle was sutured using Vicryl-coated suture thread (No.4, 74cm, Ethicon). Wound clips were used to close the skin (9mm, VWR).

### Perfusion and slice preparation

Mice were perfused transcardially at four different developmental timepoints: P7, P10, P14 and P21. A lethal dose of Sodium Pentobarbitone was intraperitoneally injected into pups, prior to perfusion with saline (0.05% in dH2O) via a peristaltic pump (Miniplus 3, Gilson) and 30G x ½ needle. Perfusate was switched to PFA 4% (in PBS) once the liver had blanched. Perfused mice were decapitated, and brains postfixed in 4% (10-12 hours). Fixed brains were sliced into 50µm coronal sections using a vibratome (Leica VT10005), which were stored in PBS (supplemented with 0.05% NaN3).

### Immunohistochemistry

Slices were thawed for 30 minutes and washed twice in PBS (5 minutes each), before permeabilization with 0.25% triton (in PBS, 4×15 minutes each). Slices were incubated for 2-hours in a blocking buffer (10% goat serum, 0.25% triton, 5% BSA). For antigen retrieval (required for VGLUT1), slices were incubated for 20 minutes (85°C) in sodium citrate buffer (10mM Sodium citrate, 0.05% Tween20, pH6.0). Slices were incubated overnight (4°C) with primary antibodies (VGLUT1, VGAT, Bassoon, RIIM1/2, Flag; Synaptic systems) at a dilution of 1:1000 in antibody solution (5% goat serum, 0.25% triton, 1% BSA). Slices were incubated for 2-hours (RT) with nanobodies at a dilution of 1:500 (FluoTag-X4 anti-GFP ATTO488, FluoTag-X4 anti-RFP AZDye568; Nanotag Biotechnologies) or secondary antibodies (Anti-Guinea Pig 680, Anti-Rabbit 647, Anti-mouse 647; Invitrogen) at a dilution of 1:1000 in antibody solution. Slices were rinsed with PBS (4×10 minutes each) and mounted with 2% agarose for super resolution (STORM) microscopy or with Mowiol for confocal microscopy.

### Confocal microscopy

Mounted slices were imaged via a Zeiss epi-fluorescent confocal microscope. Neurons were imaged using 40X or 60x oil objectives. Tiled z stacks were acquired, with 0.5µm steps and 10% overlap between tiles. Typical pixel size (X/Y was 0.12-0.16µm per pixel).

### Analysis of confocal images

Dendritic branches were traced using Simple Neurite Tracer (ImageJ Plugin^55^) and exported as swc files. ROIs were manually drawn around puncta in X-Y plane using ImageJ. Analysis of the puncta along each dendritic branch was undertaken in IgorPro (version 6.37) using custom-built scripts (Dr Guilherme Neves). Maximum Fluorescence intensity and the centroid of each ROI were measured in all 3D gaussian filtered (radius = 2 pixels) confocal slices. ROI coordinates were used to determine the nearest node within the swc trace in the XY plane, and the distance from the branch origin to the selected node was taken as the distance from the branch origin to the synapse. The maximum fluorescence in the 10 confocal sections around the z coordinate of the branch node was taken as a proxy for the synapse size. Fluorescence values for each synapse were normalized to the median of the population of all synapses of the same sign in the cell. The cumulative sum of the normalized fluorescence intensities incoming into a dendritic stretch (normalized to its length) is taken as a measure of the total synaptic input into that portion of dendrite. Correlation was assessed using Spearman’s rank correlation. Bootstrapping analysis was performed 10000 times.

Distances between synapses are measured along the dendritic shaft. Analysis of small correlation window sizes was performed in MATLAB (version 2023a). Briefly, samples of contiguous dendrite regions of varied sizes (5 to 25 microns) were taken from 10 uniformly random co-ordinates along the dendrite (random sampling was limited to two times the ratio between branch size and binning size). Correlation analysis was performed for all individual dendrite regions. This analysis was repeated 10000 times, and the average and standard deviation of the rank correlation coefficient are reported. Random placement of synapses was modelled as an exponential distribution in space, with the median distance between nearest neighbours as the lambda parameter. Synthetic dendrites were modelled with the same length and synaptic size distributions as each experimental sample, with synaptic distribution governed by the negative exponential distribution. The same number of dendrites were generated as the experimental data size and same analysis was performed in the synthetic dendrites. Synthetic branches were generated 1000 times, binning analysis was repeated 50 times for each synthetic branch, and the mean and standard deviation of the rank correlation coefficient is plotted as open circles in all figures.

### Multicolour 3D-dSTORM imaging

Brain sections were either stained solely for VGAT1 with an AF647-coupled secondary, or in some experiments for both VGAT and VGlut1 with secondary antibodies coupled to AF647 and CF680 respectively. Stained sections were mounted on 25mm coverslips coated with gelatine and placed in an AttoFluor Cell Chamber (Invitrogen). The chamber was filled with STORM buffer and sealed with another glass coverslip. STORM buffer contained final concentrations of the oxygen scavengers 100ug/mL glucose oxidase (Sigma-Aldrich G2133) and 4ug/mL catalase (Sigma-Aldrich C100), and the reducing agent 100mM b-mercaptoethylamine-HCl (Sigma-Aldrich M6500). Oxygen scavengers were prepared as a stock solution consisting of 1mg/ml glucose oxidase, 42ug/mL catalase in 20mM Tris-HCl pH7.2, 4mM TCEP, 25mM KCl, 50% glycerol. The reducing agent stock solution contained 1M b-mercaptoethylamine-HCl in deionized water, adjusted to pH8 with NaOH. Oxygen scavenger and reducing agent stocks were diluted in STORM dilution buffer consisting of 100mg/mL glucose and 10% glycerol in deionized water.

dSTORM imaging was performed using a spectral demixing SaFeRedSTORM module (Abbelight) mounted on a Nikon Eclipse Ti, equipped with a 100x/1.49 oil immersion objective. The two fluorophores AF647 and CF680 were excited with a single wavelength using a fibre-coupled 642nm laser (Errol). A long-pass dichroic beam splitter (700nm, Chroma Technology) was used to split the emission light to two ORCA-fusion sCMOS cameras (Hamamatsu). 3D-dSTORM imaging was achieved using cylindrical lenses placed in front of each camera. Prior to data acquisition, a 3D calibration was performed with Tetraspeck fluorescent beads (Invitrogen) to measure the x-y deformation obtained by the cylindrical lenses.

Image acquisition was driven by Abbelight’s NEO software. An ROI of 512×512 pixels (pixel size= 97nm) within the CA1 stratum oriens was identified using MAP2 and AF488 staining. 40,000 frames at an exposure time of 50ms were acquired using a laser power of 400mW in Hi-Lo mode. Cross correlation was used to correct for lateral drift. Single emitter events are Gaussian fit in the x-y plane using Maximum Likelihood Estimation (MLE) within the NEO software, and in the z-plane using the z-calibration data from Tetraspeck beads, thereby obtaining the 3D spatial coordinates of single fluorophores with high resolution.

NEO Analysis software was used to perform a first transformation to precisely realign detection events from both cameras using cross-correlation. A ratio of intensity between the two cameras (I1/(I1+I2)) was calculated for each detection to assign it to one of the two fluorophores, separating ratios 0-0.48 and 0.56-1. For sections stained only for VGAT, all detection ratios were included. An uncertainty maximum of 30nm and intensity maximum of 12000 were also used. Localization files (.csv) containing the 3D spatial coordinates of detections for each fluorophore were generated. Due to variation in staining efficiency, only files containing >55,000 localisations for VGAT were further analysed.

### Multicolour 3D-dSTORM analysis

Localization files were analysed using Point Clouds Analyst (PoCA)^56^, a continuation of SR-Tesseler^57^ and Coloc-Tesseler^58^. A 3D Voronoi diagram for each fluorophore was computed by creating polyhedrons centred on the localized molecules. VGAT clusters representing individual synapses were identified if they had a density at least 4 times the mean density of all localisations in the dataset. Clusters were also defined by a minimum of 50 localisations and a cutting distance of 180nm. Volumes of identified clusters were then calculated. GraphPad Prism 9 was used to assess normality and as data was non-normal, volumes were compared using a Kruskal-Wallis test with Dunn’s multiple comparisons test.

### SBFSEM

#### Sample preparation

Mice were perfused transcardially with 2% PFA in 0.1M sodium cacodylate buffer, supplemented with 2.5% glutaraldehyde. 100µm coronal sections were acquired from CA1 of the hippocampus (stratum oriens) using a vibratome (Leica, VT1000S). Tissue was incubated in potassium ferrocyanide (1.5%) at 4°C for 30 minutes and washed with dH2O for 15 minutes then incubated with aqueous thiocarbohydrazide (1%) for 4-minutes. Sections were washed again with dH2O then incubated with aqueous osmium tetroxide (2%) for 30 minutes. The sections were stained en-bloock with uranyl acetate (1%) and incubated in Walton’s Lead Aspartate for 30 minutes at 60°C, before serial dehydration in ethanol dilutions (in dH2O; 30% to 100% ethanol). Sections were infiltrated with Durcupan ACM resin (Sigma) overnight, then cured and embedded in full resin for 3 days. Sections were gold coated and mounted onto Gatan 3View aluminium pins using conductive epoxy glue (CircuitWorks).

### Image acquisition

SBFSEM imaging was undertaken at the Centre for Ultrastructural Imaging, King’s College London by Pedro Machado, using a Joel field emission scanning (JSM-7100F) electron microscope with a 3View 2XP system (Gatan) with high vacuum pressure at 2.5kV and a 8000X20000 scan rate covering a total area of 40 × 100 mm with a pixel size of 5 nm. 662 slices were acquired at a step size of 50 nm resulting in a total depth of 33.1µm.

### SBFSEM analysis

Stacks were semi-automated aligned using the linear stack alignment with SIFT ImageJ plugin. Dendrites were segmented using the ImageJ plugin TrakEM2 (59) then reconstructed in 3D and analysed in Blender (version 2.91.2) with the Neuromorph toolset plugin (60). Further analysis was undertaken using Igor Pro (version 6.37) with custom built scripts. GraphPad Prism (version 9.1.1) was used for statistical analysis and graph production. Synapse size was estimated by measuring the post-synaptic density (PSD) surface area of each synapse. Location of synapses was estimated using the centerline function of Neuromorph. The analysis of cumulative synaptic input used was modelled on the measures obtained from the confocal images, replacing the normalized fluorescence intensities with the PSD surface areas.

### Statistical Analysis

All electrophysiology data was analysed and plotted using a custom-made Python script. Graphpad Prism (version 9.1.1) was used for statistical analysis of the confocal, dSTORM and SBFSEM data. Correlations were assessed using Spearman’s rank correlation, and bootstrapping analysis was performed 10000 times, and the average and standard deviation of the rank correlation coefficient are reported. GraphPad Prism (version 9.1.1) was used to assess normality of dSTORM data and as data was non-normal, volumes were compared using a Kruskal-Wallis test with Dunn’s multiple comparisons test.

## Supplementary Figures

**Figure S1:**
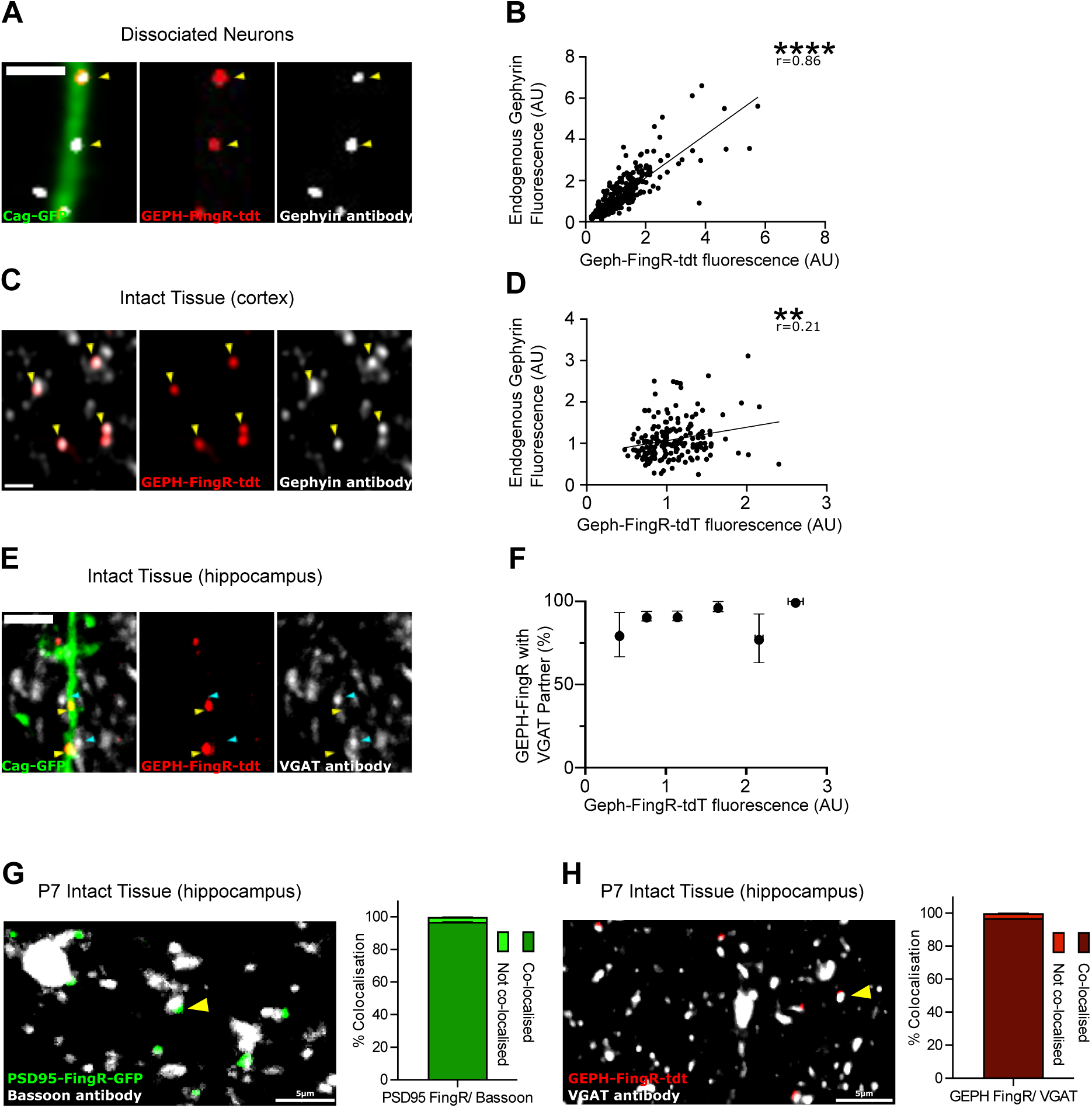
PSD95-FingR-GFP and GEPH-FingR-tdt label bona fide synapses and report synapse strength. (A) An example image of a dissociated hippocampal neuron co-expressing GEPH-FingR-tdt and CAG-GFP and immunolabelled for endogenous gephyrin. (B) A significant positive correlation was observed between the amplified GEPH-FingR-tdt fluorescence intensity and the intensity of signal from anti-gephyrin antibody (n=336, r=0.86, P<0.0001). (C) An example image of a cortical pyramidal neuron expressing GEPH-FingR-tdt and CAG-GFP via in utero electroporation and subsequent immunolabelling for endogenous gephyrin. (D) Normalised fluorescence intensity of GEPH-FingR-tdt puncta were plotted against the fluorescence intensity of endogenous gephyrin antibody puncta and a significant positive correlation was observed (n=138, r=0.2126, P=0.0039). (E) An example image of hippocampal pyramidal neuron expressing GEPH-FingR-tdt and CAG-GFP via in utero electroporation, with subsequent immunolabelling for VGAT. (F) GEPH-FingR-tdt puncta were binned based on fluorescence intensity, and plotted against the percentage that were colocalised with a VGAT partner. Across all bins, 92% GEPH-FingR-tdt puncta had a VGAT partner (n=239). (G) Example image of hippocampal neurons expressing PSD95-FingR-PSD via in utero electroporation in brains fixed at P7, with subsequent immunolabelling for Bassoon. 97% PSD95-FingR-GFP puncta colocalised with Bassoon puncta at P7. (H) Example image of hippocampal neurons expressing GEPH-FingR-tdt via in utero electroporation in brains fixed at P7, with subsequent immunolabelling for VGAT. 97% GEPH-FingR-GFP puncta colocalised with VGAT puncta at P7. (A, C, F) Scale bars represent 1µm. (G, H) Scale bars represent 5µm.

**Figure S2:**
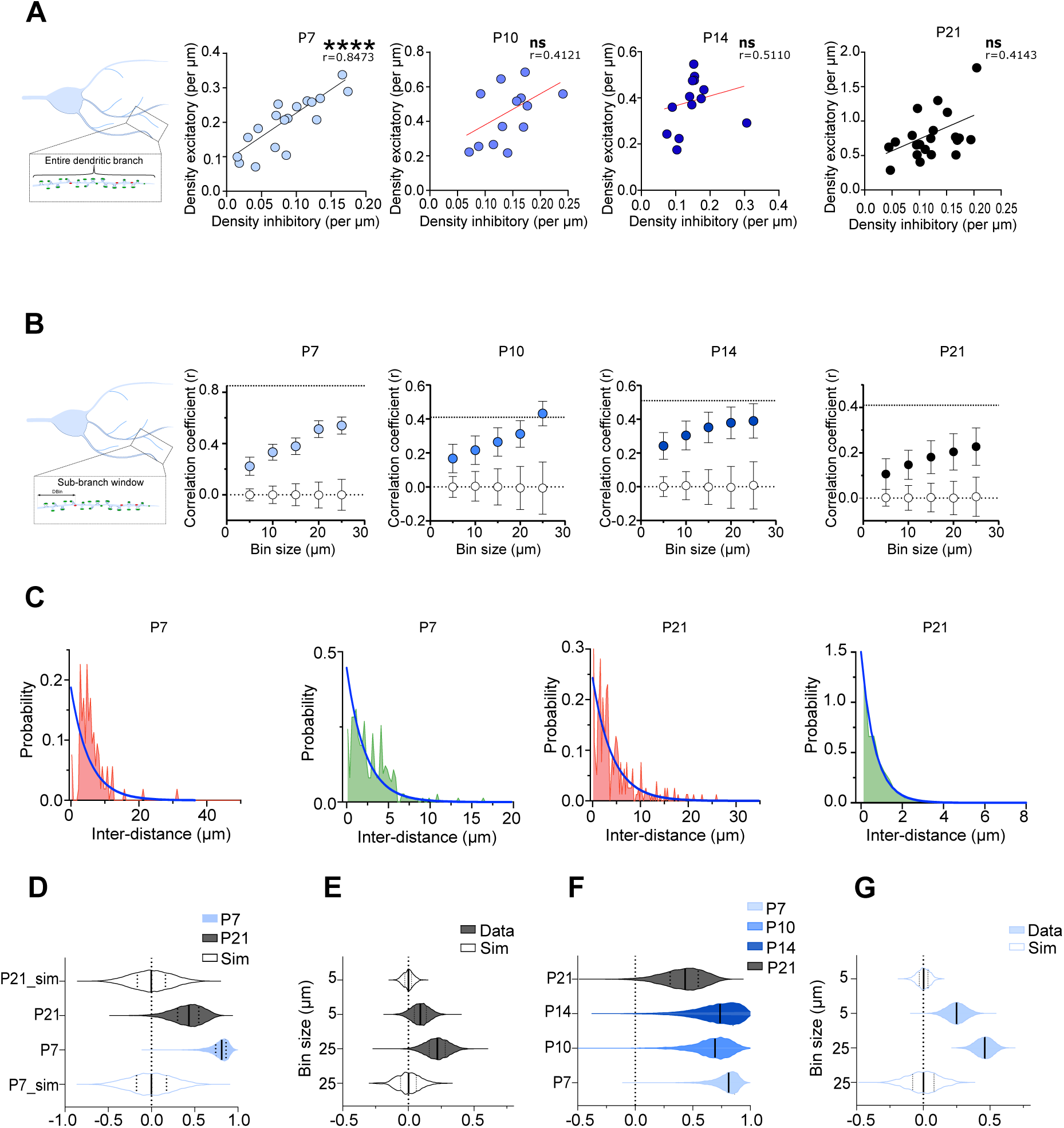
The density of excitatory synapses and the density of inhibitory synapses are correlated at a sub-branch level throughout development. (A) Schematic showing how the balance between synapse density was assessed along entire dendritic branches, and graphs showing the density of excitatory synapses as a function of inhibitory synapses at P7, P10, P14 and P21. A significant positive correlation was observed at P7 (r=0.8473, P<0.0001, n=18 dendrites from 6 cells). No correlation was identified at P10 (r=0.4121, P=0.1635, n=13 dendrites from 3 cells), P14 (r=0.5110, P=0.0776, n=13 dendrites from 2 cells), or P21 (r=0.4143, P=0.0619, n=21 dendrites from 7 cells). (B) Schematic showing how the balance between synapse densities was assessed within sub-branch windows (DBin) of different lengths, and graphs showing the correlation coefficient as a function of Bin Size throughout development (P7 to P21). Correlation coefficient increased with bin size for imaged (close circles) but not simulated (open) branches with random synaptic placement at all analysed ages. Dotted lines represent zero correlation, and the coefficient calculated using the entire branches as shown in Figure S 2A. (C) Histogram of Nearest Neighbour distances (NN) (normalized to the probability density function) for inhibitory (red) and excitatory (green) synapses compared with the probability density function of the exponential distribution with mean equal to the median nearest neighbour distance of the population (blue line). The exponential distribution closely matches only the distribution of NNs for P21 excitatory synapses. (D) Violin plot of the Spearman rho population data from either bootstrapping analysis of observed data (10000 resampling, filled plots, median and 25 and 75 quartiles indicated by lines, datasets used-P7-blue – from Fig 2H and P21-gray – from Fig. 1E), or random generation of simulated branches (10000 repeats, open plots). (E) Violin plot of the population of Spearman rho data generated with the random analysis of dendritic stretches of different sizes (bins) from the P21 confocal analysis summarised in Fig 1F. Analysis of 5 and 25 micron sized bins is illustrated. Open plots represent similar analysis on simulated branches. (F) Violin plot of the distribution of spearman rho coefficients obtained by bootstrapping analysis across development. (G) Violin plot of the population of Spearman rho data generated with the random analysis of dendritic stretches of different sizes (bins) from the P7 confocal analysis summarised in Fig 2J. Analysis of 5 and 25 micron sized bins is illustrated. Open plots represent similar analysis on simulated branches.

**Figure S3:**
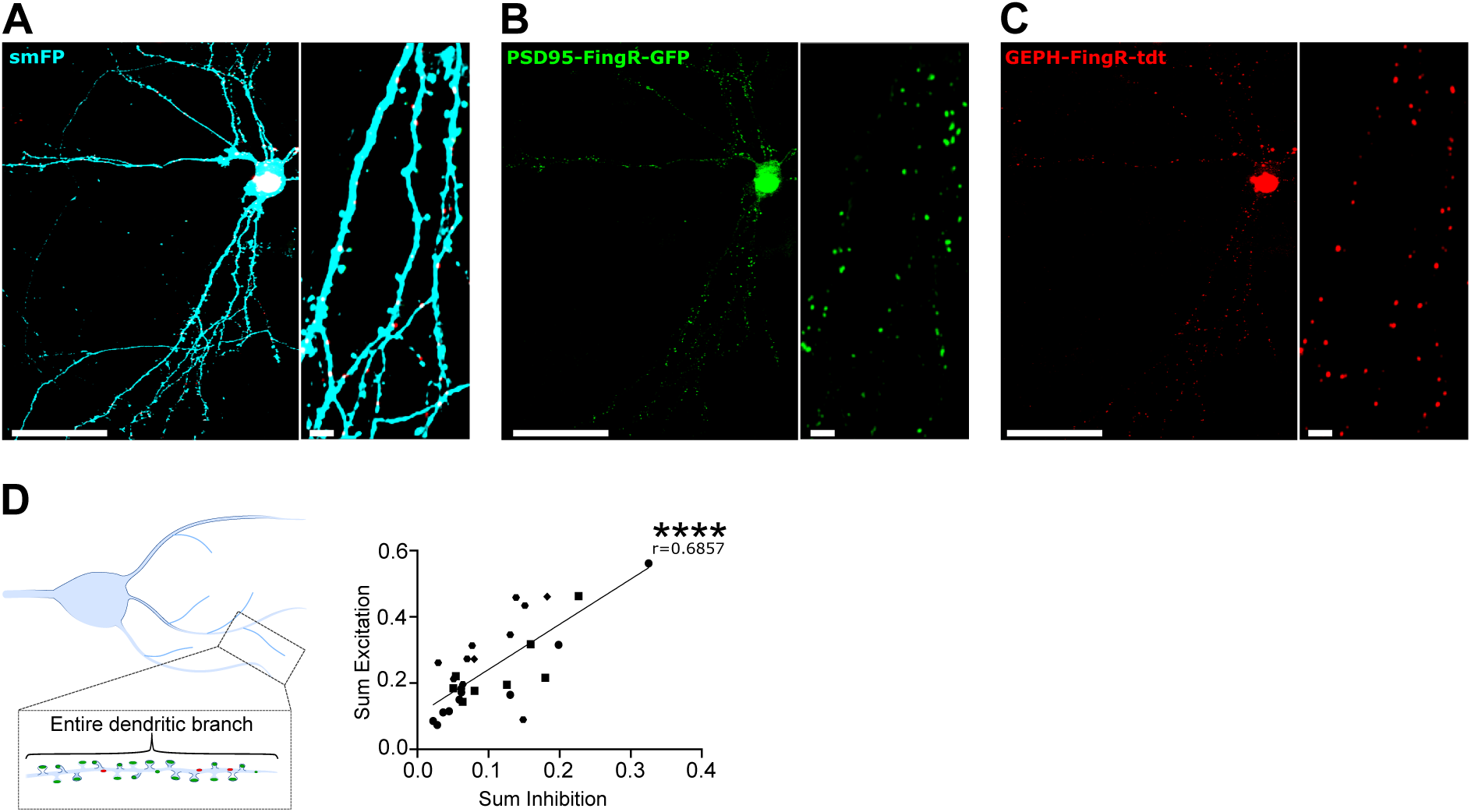
Excitation and inhibition are balanced at a subcellular level in dissociated hippocampal neurons. (A-C) An example image of a neuron at 21DIV expressing (A) smFP, (B) PSD95-FingR-GFP and (C) GEPH-FingR-tdt to label neuronal structure, excitatory synapses and inhibitory synapses, respectively. (D) Schematic showing how the balance between the sum of excitation and the sum of inhibition was assessed along entire dendritic branches. A graph showing the sum of excitation as a function of the sum of inhibition, and a significant positive correlation was observed (r=0.6857, P<0.0001, n=29 dendrites from 4 cells). Scale bars represent 50µm (larger images) and 5µm (magnified images).

